# A novel approach for the quantification of single-cell adhesion dynamics from microscopy images

**DOI:** 10.1101/2024.10.08.616409

**Authors:** Marilisa Cortesi, Jingjing Li, Dongli Liu, Tianruo Guo, Socrates Dokos, Kristina Warton, Caroline E. Ford

## Abstract

**Background:** Cell adhesion, that is the ability to attach to a given substrate, is a key property of cancer cells, as it relates to their potential for dissemination and metastasis. The *in vitro* assays used to measure it, however, are characterized by several drawbacks, including low temporal resolution and limited procedural standardisation which reduce their usefulness and accuracy.

**Results:** In this work, we propose an alternative analytical approach, based on live-cell imaging data, that yields comprehensive information on cell adhesion dynamics at the single-cell level. It relies on a segmentation routine, to identify the pixels belonging to each cell from time-lapse microscopy images acquired during the adhesion process. A tracking algorithm then enables the study of individual cell adhesion dynamics over time. The increased resolution afforded by this method was instrumental for the identification of cell division prior to attachment and the co-existence of markedly different proliferation rates across the culture, previously unidentified patterns of behaviour in the adhesion process. Finally, we generalize our method by substituting the segmentation algorithm of the instrument used to acquire the images, with a custom-made one, showing that this approach can be integrated within routine laboratory analytical procedures and does not necessarily require high-performance microscopy and imaging setups.

**Conclusions:** Our new analytical approach improves the *in vitro* quantification of cell adhesion, enabling the study of this process with high temporal resolution and increased level of detail. The extension of the analysis to the single-cell level, additionally, uncovered the role of population variability and proliferation in this process. The simple and cost-effective procedure here described enables the accurate characterisation of cell adhesion. Beside improving our understanding of adhesion dynamics, its results could support the development of treatments targeting the ability of cancer cells to adhere to surrounding tissues.

## Background

The ability of cancer cells to adhere to a given substrate is a key property, which can provide insight into their metastatic potential as swift and robust adhesion to the healthy tissue is a key step toward its colonisation. The *in vitro* study of this phenomenon largely consists in (i) seeding a known number of cells on the substrate of interest, (ii) waiting a set amount of time, (iii) washing away the non-attached cells and (iv) quantifying the density of the adherent population, generally through an absorbance measurement (1, 2). While procedurally simple and largely accessible, as it does not require specific instrumentation, this assay is characterised by many drawbacks. These include binary classification in attached/detached, rather than the identification of the different phases of adhesion, and restriction of the analysis to population level, that eliminates intercellular variability, a fundamental element of cancer progression and metastasis initiation (3). The results are also dependent on the intensity of the washing step, which is difficult to standardise and could potentially vary between samples and operators. While alternative methods are available (4), and some of them enable the evaluation of single cell adhesion strength (5), they generally require specific instrumentation, and the experimental conditions tend to be quite far from *in vivo*-like culturing setups.

Live-cell imaging analysis offers a potential solution. Indeed, different adhesion stages result in morphological changes in the cells (6) which can be identified with an appropriate segmentation and classification routine. An advantage of these setups is that they maintain cells in standard culturing conditions, thus minimising the effect of the measurement on the biological process itself (7–9).

In this study, we evaluated the feasibility of this approach by using an IncuCyte S3 Live Cell Analysis system to acquire images of two different high grade serous ovarian cancer (OC) cell lines (PEO1 and PEO4) as they adhered to the bottom of a multi-well plate. While this method is potentially applicable to any cell type, OC is a particularly interesting application as metastasis formation is common, more than 75% of patients present at diagnosis with metastatic disease (10), and bears a strong association with poor outcome and development of drug resistance (11, 12), Adhesion is also particularly important for OC as it disseminates almost exclusively through the transcoelomic route, a process by which cancer cells spread across a body cavity, the peritoneal space in this case, rather than infiltrating the circulatory system (13, 14).

The selection of PEO1 and PEO4 cells is also interesting, as they were derived from the same patient, but represent different stages of OC progression, first recurrence (PEO1) and the development of treatment resistance (PEO4) (15). These feature makes them highly comparable, while allowing for a range of behaviours to be explored.

Our results show that cell adhesion is a highly heterogeneous and dynamic process and provide a novel approach to quantify it at high resolution and in an operator independent, highly automated manner.

## Results

Adhesion was monitored in PEO1 and PEO4 cells by seeding different concentrations of these cells in standard 96-wells plates immediately prior to their positioning within an IncuCyte S3 Live Cell Analysis system. Images of the same area of the culture were acquired for 20 h with a temporal resolution of 30 minutes. Analysis of these images (further details in the methods) enabled the determination of each cell’s adhesion status.

### Adhesion at the population-level is dynamic and cell line-dependent

The same qualitative behaviour was observed, at the population level for all conditions: the fraction of detached cells decreased as that of partially attached, and fully attached cells increased (Figure 1). However, some cell line specific dynamics were observed. PEO1 cells, adhered more quickly, with the population of detached cells dropping below 20% before the end of the experiment and the rest of the population uniformly split between partially and fully attached. PEO4 cells, on the other hand, remained mostly in the intermediate partially attached state, with only about 20% of the cells attaining complete adhesion after 20 h.

**Figure 1:**
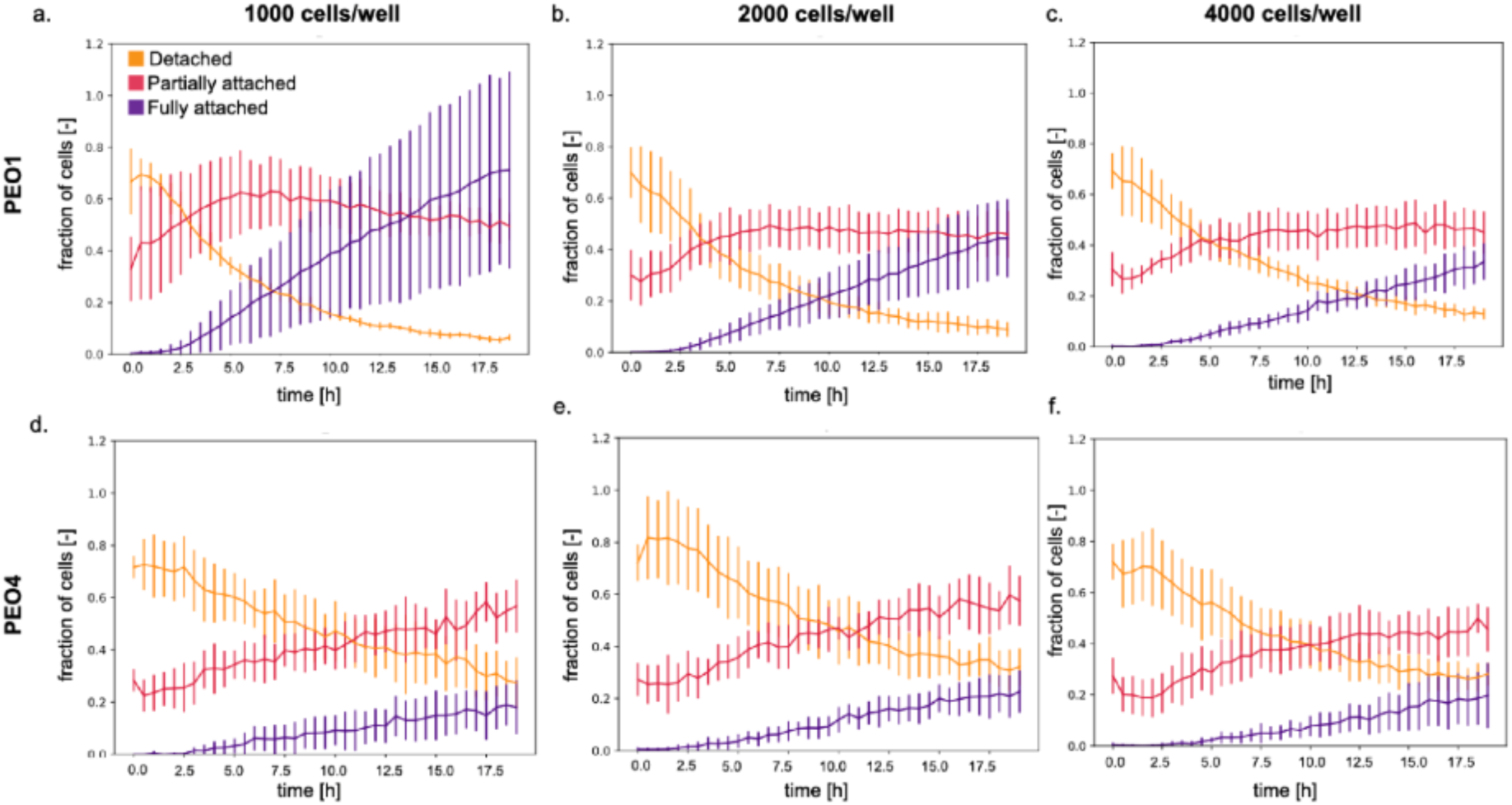
Global adhesion analysis showing the prevalence of each adhesion class over time. a. Average and standard deviation of the fraction of cells for each adhesion status (detached in orange, partially attached in red and fully attached in purple) as measured for PEO1 cells with an initial cell density of 1000 cells/well (3 wells and 3 images/well). b. and c. are the same as a. but with starting populations of 2000 and 4000 cells/well. d., e. and f. are the same as a., b., and c. but for PEO4 cells.

Changing the initial cell density (1000, 2000 or 4000 cells/well) affected the behaviour of PEO1 cells, with a reduction in adhesion speed proportional to the population size. Indeed, the time point at which the fully attached population becomes more prevalent than the detached one is 7 h for the 1000 cells/well and 13 h for 4000 cells/well. This effect is not present for PEO4 cells, where the difference in prevalence between the different classes is largely conserved when changing the seeding density. This is coherent with PEO4 cells being derived from a more advanced, aggressive disease stage. Adhesion assays conducted using the standard approach by Ritch et al. yielded comparable results: PEO1 cells adhered more and with a faster dynamic when compared with PEO4 cells (16). The time courses presented here, however, are characterized by a higher temporal resolution (30 minutes instead of a few hours) and enable the identification of the different adhesion phases.

### Distinct adhesion dynamics identified by single-cell level adhesion analysis

The quantification of adhesion through the segmentation and analysis of live cell imaging data also supports the characterisation of adhesion at single cell level. In particular, cells in consecutive images were matched through a Bayesian cell tracking algorithm (btrack) relying on a probabilistic model of cell movement, spatial information and cell appearance (17). Pairing this information with the corresponding adhesion status led to the identification of nine different adhesion dynamics (Table 1).

**Table 1:**
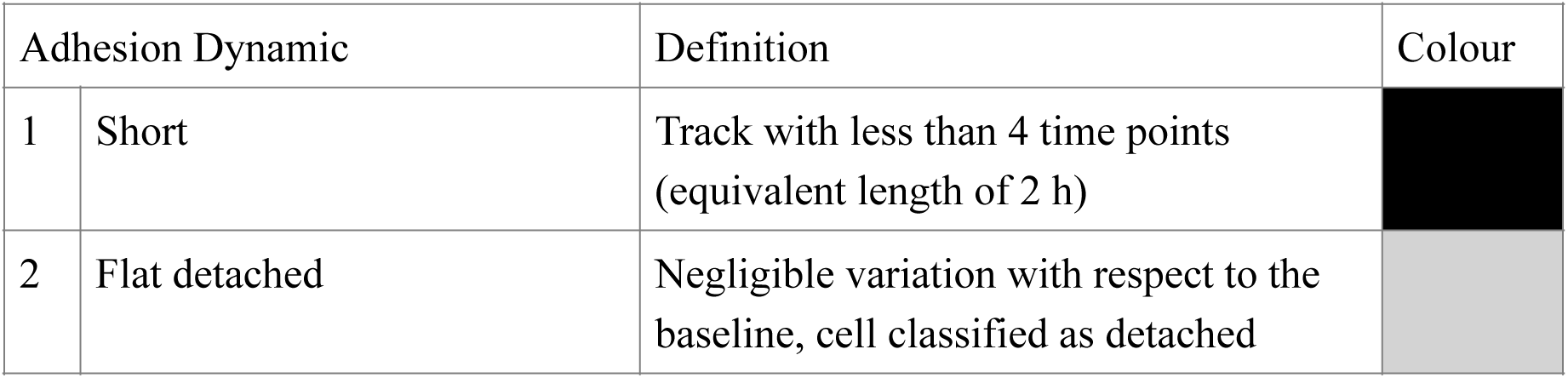
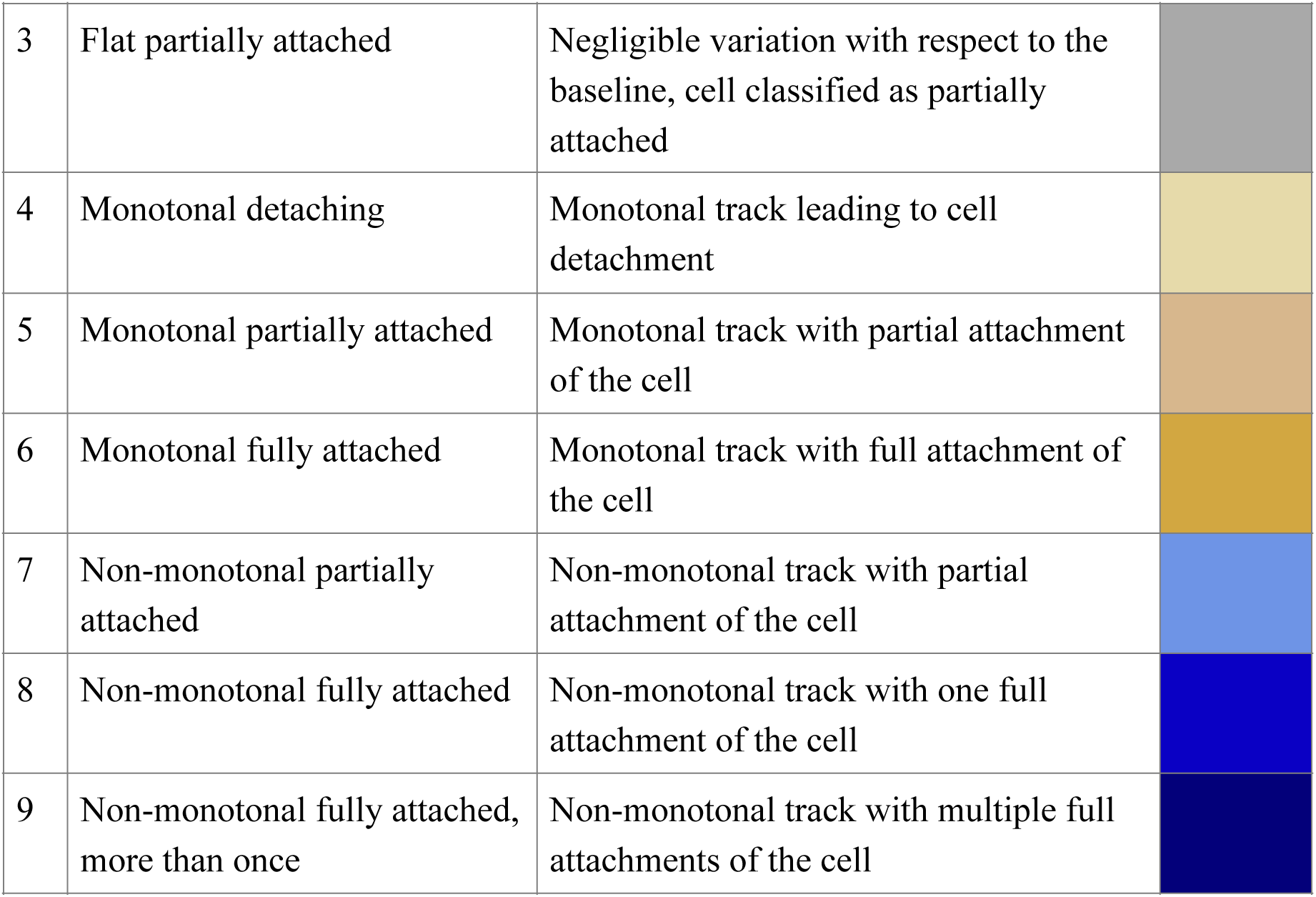
Summary of the different adhesion dynamics, their definition and associated number and colour.

| Adhesion Dynamic |  | Definition | Colour |
| --- | --- | --- | --- |
| 1 | Short | Track with less than 4 time points (equivalent length of 2 h) |  |
| 2 | Flat detached | Negligible variation with respect to the baseline, cell classified as detached |  |
| 3 | Flat partially attached | Negligible variation with respect to the baseline, cell classified as partially attached |  |
| 4 | Monotonal detaching | Monotonal track leading to cell detachment |  |
| 5 | Monotonal partially attached | Monotonal track with partial attachment of the cell |  |
| 6 | Monotonal fully attached | Monotonal track with full attachment of the cell |  |
| 7 | Non-monotonal partially attached | Non-monotonal track with partial attachment of the cell |  |
| 8 | Non-monotonal fully attached | Non-monotonal track with one full attachment of the cell |  |
| 9 | Non-monotonal fully attached, more than once | Non-monotonal track with multiple full attachments of the cell |  |

These vary greatly in terms of length, complexity and prevalence within the population (Figure 2). For both cell lines, short tracks like the ones shown in Figure 2a were the most common (86 and 89% for PEO1 and PEO4 respectively). The main reason for their prevalence is that they combine several different situations, including cells that were not alive at the time of seeding, cells that exited the field of view or newly divided cells at the end of the experiment. They were however excluded from most of the analysis that follows as they would have required the use of separate methods, due to their short length and their adhesion status was mostly constant throughout the experiment.

**Figure 2:**
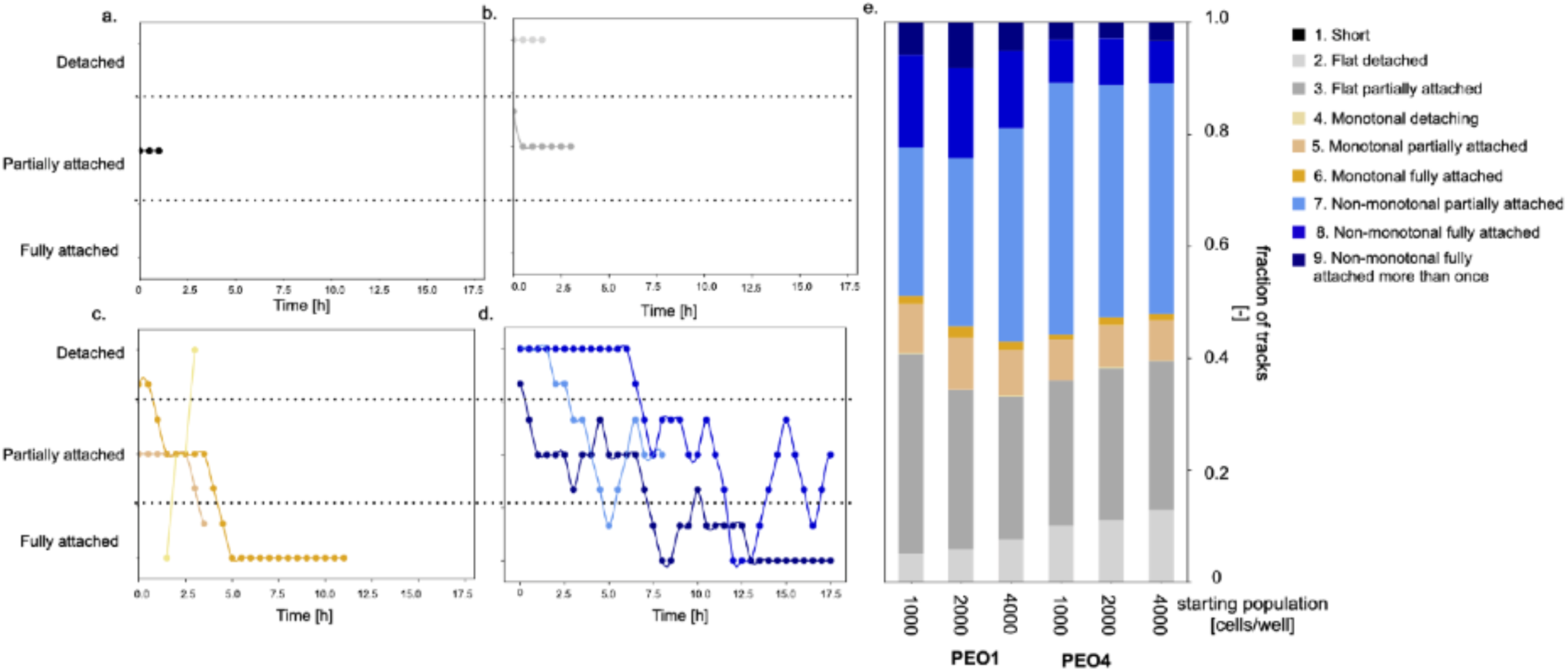
Analysis of the adhesion dynamics of individual cells. a. Representative example of a short track, i.e., a track shorter than 2 h. b. Example of flat tracks. These are longer than short tracks but are also defined by a small dynamic range. Flat tracks can be either detached or partially attached. c. Monotonal tracks maintain the same direction of change. They can be upward (detaching) or downward (attaching). The latter has been divided between attaining full attachment (monotonal fully attached) or remaining at the partially attached stage (monotonal partially attached). d. non-monotonal adhesion tracks can also attach completely (non-monotonal fully attached) or stop at the partially attached stage (non-monotonal partially attached). In some cases, full attachment was observed multiple times for the same cell (non-monotonal fully attached more than once). d. Breakdown of the prevalence of each trace type for each considered condition (PEO1 and PEO4 cells, 1000, 2000 and 4000 cells/well). Short tracks were excluded in the making of this plot and each dot in panels a-d represents a measure.

Flat partially attached tracks (representative example shown in Figure 2b) were also common (Figure 2e, Table 2), together with the non-monotonal behaviour leading to a partial attachment of the cell (light blue in Figure 2d, e and Table 2). Monotonal tracks (representative examples in Figure 2c) were overall the least likely (Figure 2e, Table 2). Attainment of full attachment was the most common for PEO1 cells (on average 23% of the non-short tracks vs 12% for PEO4 cells). A statistical analysis of the prevalence of each adhesion dynamic has been conducted to show how the median likelihood of each track type changes with cell line and seeding density (Supplementary Figure 1). Differences between cell lines are the most common, even though variation among seeding densities was also observed (e.g., Flat partially attached tracks for PEO1 cells Supplementary Figure 1b).

**Table 2:**
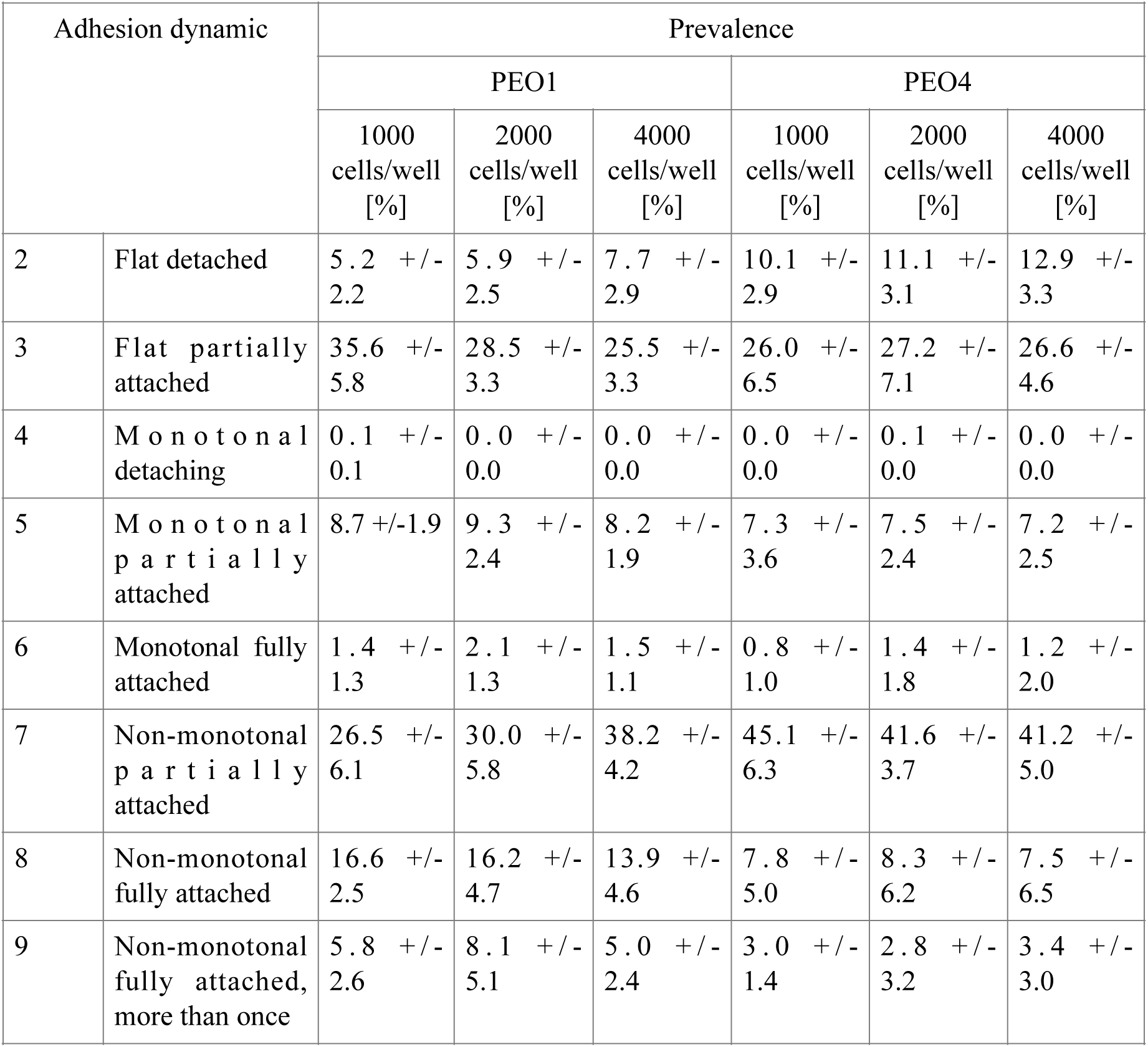
Percentage prevalence of each adhesion dynamic organised by cell line and initial cell density. These values were calculated after the exclusion of the short tracks. Statistical analysis for these data is provided in Supplementary Figure 1.

### The length of the adhesion track is propor3onal to its complexity

An analysis of the tracks’ length (Figure 3) revealed some interesting differences both between adhesion dynamics and cell lines. Track complexity (e.g., non-monotonicity, attainment of full adhesion) was directly correlated with track length, with the “non-monotonal fully attached, more than once” behaviour associated with a median length of at least 12.5 h. (Figure 3a). A high variability in track length was however recovered for most track types (Figure 3). This was particularly evident for PEO4 cells, where the interquartile range spanned more than 10 h for most conditions (Figure 3b, d, f).

**Figure 3:**
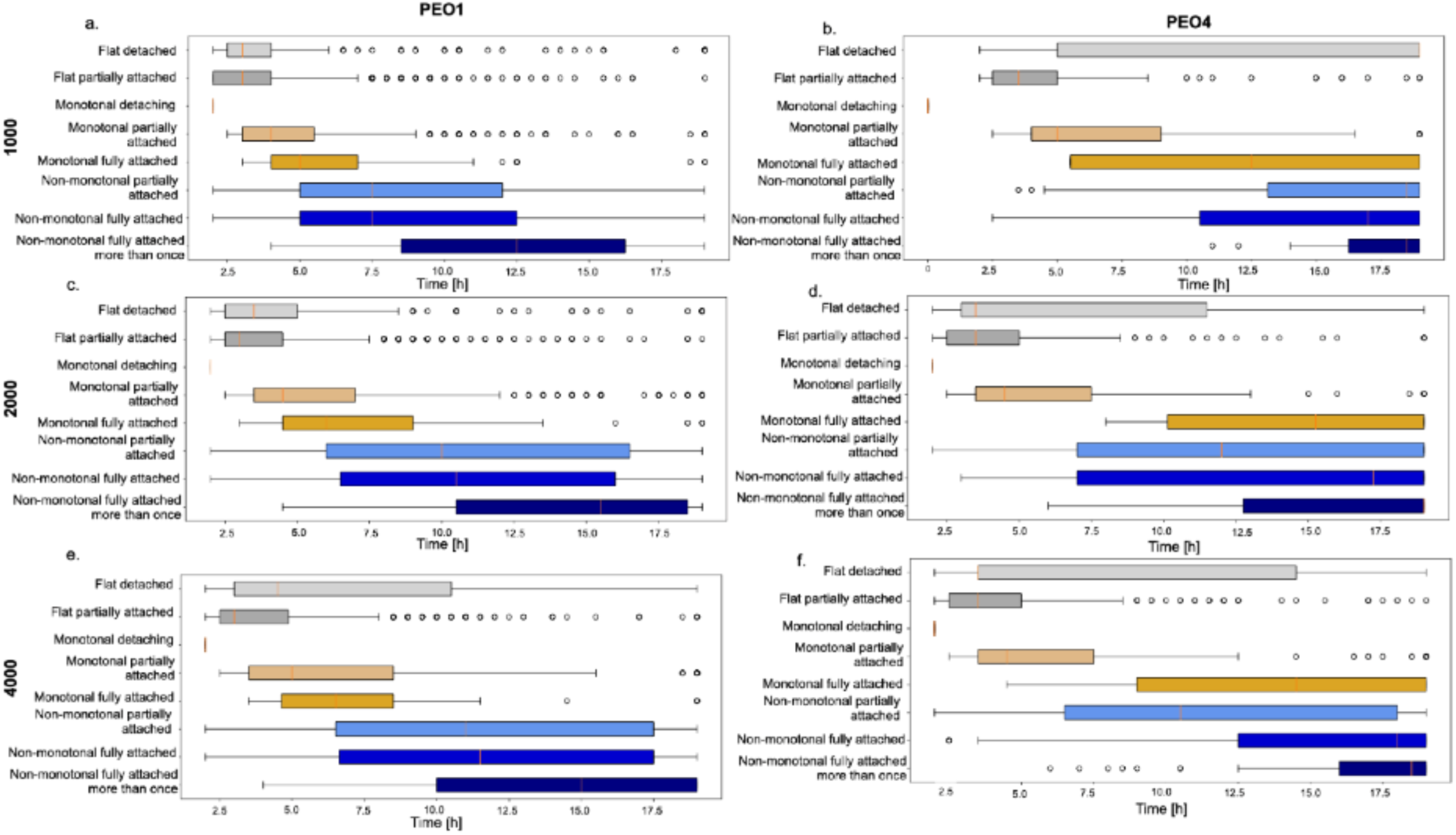
Study of the distribution of track length. Each panel reports the result for each cell line and initial cell density. The boxplots describe the distribution of track length for each adhesion dynamic (identified by the labels on the y-axis and the colours). Orange bars mark the median of the distributions, while the edges of the coloured boxes are the interquartile range. Short tracks (tracks with length less than 2 h) were excluded.

### Cell doubling occurs early, and its probability depends on the cell’s adhesion dynamic

One of the main features of btrack is the lineage tree reconstruction which enabled the identification of cell doubling and kept track of each cell’s parent and offspring. Despite the average doubling time for both PEO1 and PEO4 cells being longer than the duration of our experiment (at least 36 h (18)), we detected several cell divisions during our experiments. Figure 4a shows the growth ratio, defined as the ratio between final and initial number of cells. PEO1s have median growth ratio of about 1, corresponding to the maintenance of a stable population. A non-negligible variability among the replicates was however observed, together with trend toward lower growth when a higher cell density was considered. This dependence of the proliferation rate on the population size was further confirmed by the fraction of cells that were observed dividing during the experiment (Figure 4b). Indeed, the number of proliferating PEO1 cells more than doubled when the initial population density was lowered from 4000 to 1000 cells/well.

**Figure 4:**
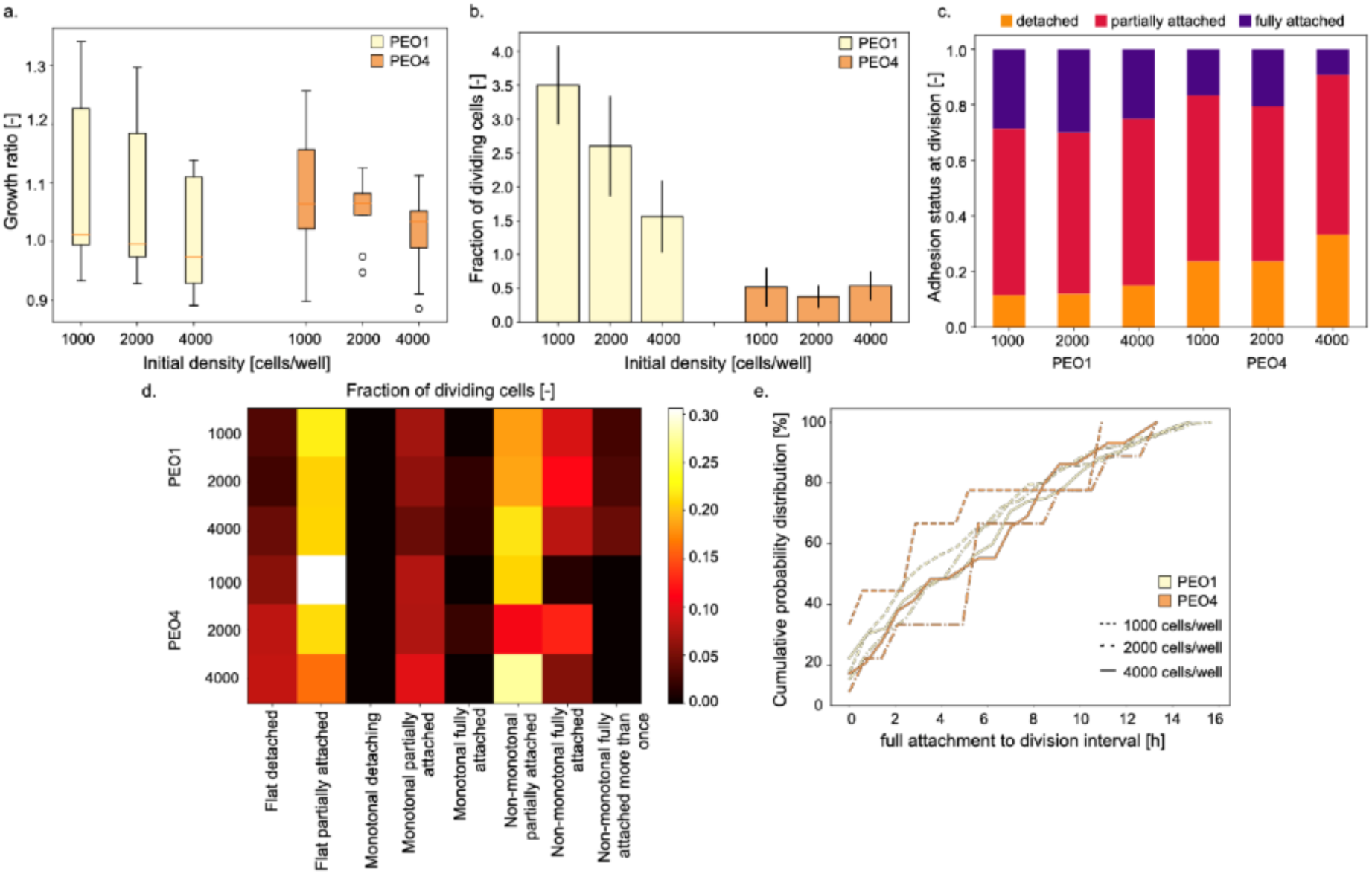
Analysis of cell doubling (conducted excluding short tracks). a. Distribution of growth ratios as a function of cell line and initial cell density. b. Fraction of cells for which division was recorded organised by cell line and initial density. c. Prevalence of each adhesion status (orange detached, red partially attached and purple fully attached) recorded at cell division. d. Fraction of dividing cells by adhesion dynamic. e. Cumulative distribution of the time interval between attainment of full attachment and cell division. The cumulative distribution represents the likelihood of the variable on the x axis to be less than or equal to the reported values.

PEO4 cells, on the other hand, showed a slightly higher median growth ratio (about 1.1) and a reduced dependency on the initial population size (Figure 4a, b). Another important difference was the reduction in the fraction of proliferating cells observed for this cell line which dropped by about an order of magnitude (Figure 4b).

Division was most likely to occur when cells were detached or partially attached (Figure 4c). Figure 4 d. reports the breakdown of the fraction of dividing cells by adhesion dynamic. Interestingly, full attachment was not required for doubling to occur. Indeed, for both PEO1s and PEO4s, most of the cells that were observed dividing were classified as “flat partially attached” and “non-monotonal partially attached”. Full attachment was generally associated with low probability of division, especially following a monotonal track. When cells attaining full attachment divided, they were likely to do it soon after completing adhesion (Figure 4 e). Indeed, the cumulative distribution of the interval between full attachment to the substrate and cell division reached 50% between 4 and 6 hours for all conditions.

We then focussed our attention on the daughter cells, that is the cells that were recorded by btrack to have a parent id different from their own. Figure 5 shows the distributions of track start times for these cells, divided by cell line and initial population density. Each dot represents a cell, color-coded according to its adhesion dynamic. This plot clearly shows the difference in number of division events recorded for each condition, even though the median start time was about 13 h in all cases. More complex tracks are generally associated with an earlier start time, but full adhesion was still observed in cells whose tracking started less than 5 h from the end of the experiment.

**Figure 5:**
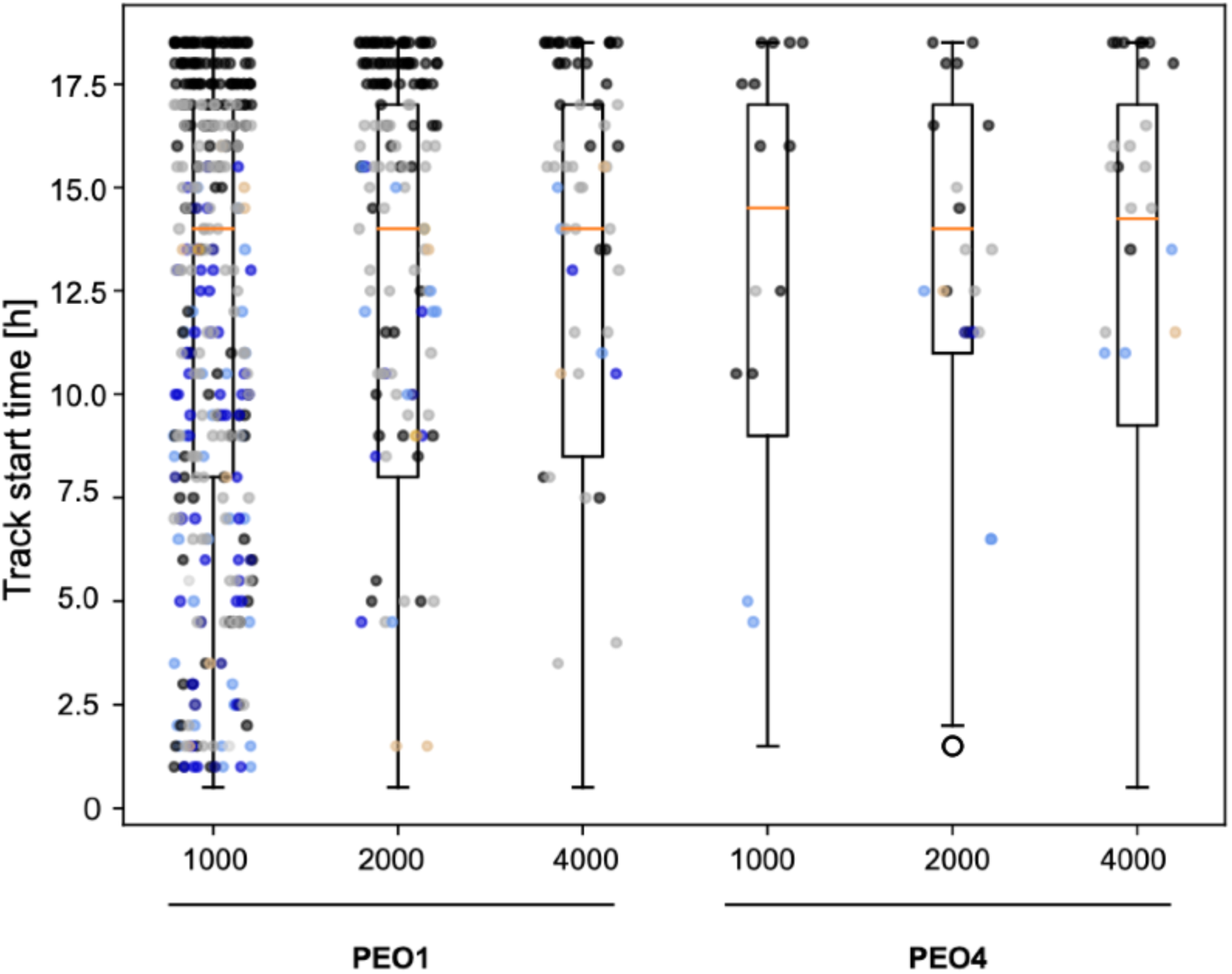
Start times distributions for the daughter cells, that is the ones that were not present when the experiment started. Each dot represents a cell and is color-coded according to the corresponding adhesion dynamic.

### Substituting the segmentation algorithm does not affect the results of the adhesion analysis

As highly specialised instruments such as the IncuCyte S3 Live Cell Analysis system are unlikely to be widely available, we decided to repeat the same analysis with a custom-made segmentation software (18). This would extend the applicability of our method to other live cell monitoring systems and images acquired with standard microscopy setups. This analysis was undertaken only for two initial cell densities (2000 and 4000 cells/well). Figure 6 shows the comparison between the results obtained with the algorithm in (18) and the ones presented in the previous sections. In a. the Jaccard index (Eq 1) of the two segmentations is shown. This is a similarity metric that evaluates the fraction of shared foreground pixels between two segmented images (A and B) by dividing their intersection, or area of overlap (|A∩B|) by the total foreground area (e.g., their union, |AUB|).

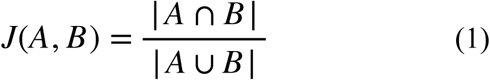

**Figure 6:**
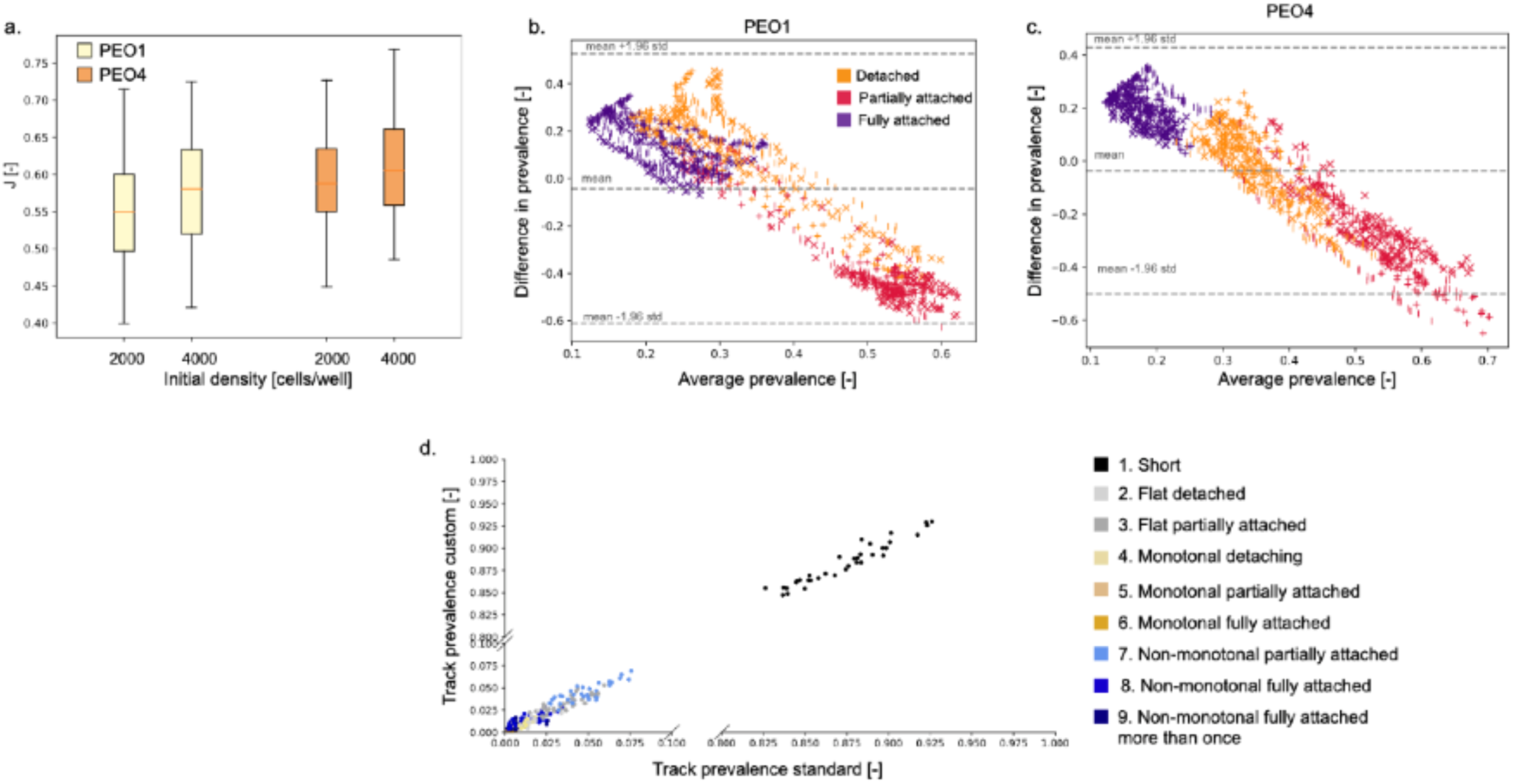
Comparison between the results obtained with the two segmentation methods. a. Jaccard index (Eq. 1) quantifying the superimposition between the two segmentations of the same image. b. Bland Altman plot comparing the two segmentation algorithms for PEO1 cells. Each point represents the prevalence of one adhesion class at a specific timepoint (initial cell densities and replicates have been kept separate). Different colours identify the corresponding adhesion stages (orange: detached, red: partially attached and purple fully attached), while markers were used to separate the initial cell densities (x: 2000 cells/well, |: 4000 cells/well). c. Same as b. but for PEO4 cells. d. Correlation between the prevalence of each track type as measured by the standard segmentation algorithm (x axis) or the custom approach (y axis). Colours identify the different track types.

A comparable distribution of Jaccard indices was observed for all tested conditions (Figure 6a.), confirming that cell shape and density did not affect the quality of the segmentation. The analysis of the global adhesion dynamics was also equivalent (Figure 6b.c. and Supplementary Figure 2). This was demonstrated through the Bland Altman plot, that compares the difference between the results of two methods to their average. As most points (99.9 and 97.4% for PEO1 and PEO4 respectively) are within the 95% confidence interval (Figure 6 b, c), the global adhesion analysis obtained with custom segmentation can be considered equivalent to the one shown in Figure 1 (19, 20). This is also supported by the global adhesion dynamics obtained with the custom segmentation (Supplementary Figure 2). Comparing these graphs to the ones in Figure 1, the same trends and overall behaviour is observed. The main difference was the initial prevalence of partially attached PEO4. Indeed, they are about as numerous as the detached ones in Supplementary Figure 2, while in Figure 1 they are about half as likely. This was probably connected with the classification procedure and the thresholds used to distinguish the different adhesion states. These values were empirically chosen for the segmentations obtained with the Incucyte proprietary software but were also applied when the custom algorithm was used. This choice simplifies the comparison between the two methods and was justified by the high similarity between the two segmentations (Figure 6a) but might lead to a slightly sub-optimal classification of the adhesion stages for the custom algorithm.

The prevalence of the different adhesion dynamics was also highly correlated (R^2^ = 0.999, Figure 6d., Supplementary Figure 3). This result confirms that the equivalence of the two segmentation methods extends to the single cell level analysis. Indeed, very similar tracking results were obtained, both in terms of track type prevalence (Supplementary Figure 3 and Figure 2d) and track length (Supplementary Figure 4 and Figure 3).

### The use of a custom segmentation algorithm does not affect the analysis of early cell doubling

Figure 7a shows the growth rate distribution obtained with the custom algorithm. When compared with Figure 4a, the median population growth was slightly higher (about 1.2 for all conditions) and the effect of initial cell density was less noticeable. The fraction of doubling cells, however, was highly consistent between the two methods (Figure 7 b and Figure 4 b). PEO1 cells were associated with a higher fraction of dividing cells and there was a trend toward reduced proliferation when a higher initial cell density is considered. PEO4 cells, on the other hand, maintained a similar fraction of doubling cells independently of the initial population. Breaking down the fraction of dividing cells according to their adhesion dynamic also yielded similar results. Beside directly comparing Figure 7c to Figure 4c we also calculated the error distribution for each track type (Figure 7d). While the distribution variability changed between the different adhesion dynamics the error was below 10% in all cases.

**Figure 7:**
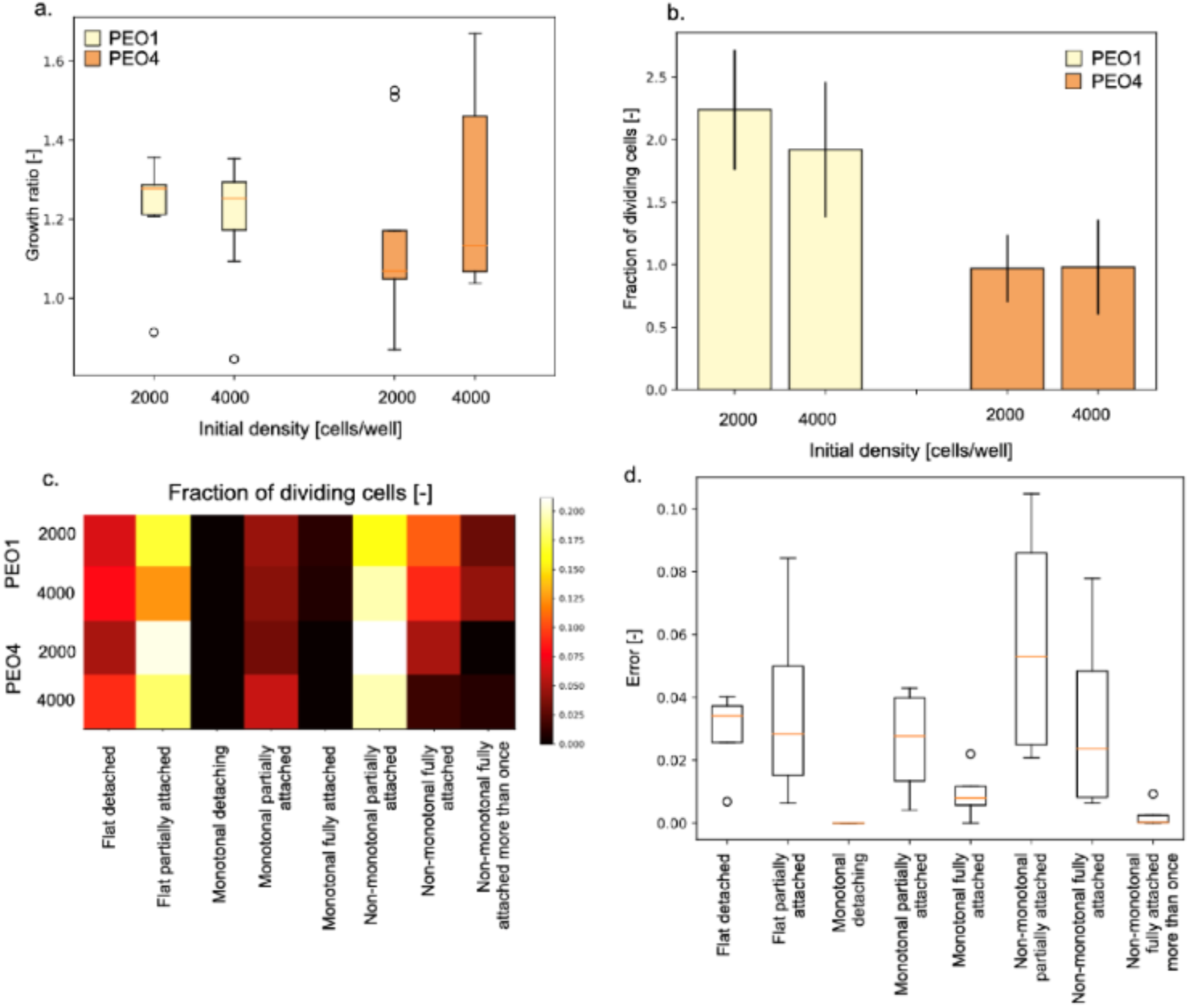
Analysis of cell doubling using the custom segmentation algorithm. a. Growth ratio distribution measured for each cell line and starting condition. b. Average (+/- standard deviation) of the fraction of dividing cells. Each bar comprises data from 3 wells (3 image stacks/well). c. Heatmap showing the fraction of dividing cells organised by track type. d. Error between the fraction of dividing cells in Figure 5 c and Figure 8 c.

The start-time distribution for the daughter cells was also largely conserved between the two segmentation algorithms (Figure 8 and Figure 5). The median start time was just under 13 h and there was a direct correlation between track complexity and start time. While there was some variation in the number and distribution of recorded division events (colour coded dots in Figures 8 and 5), the same conclusions can be derived from both figures.

**Figure 8:**
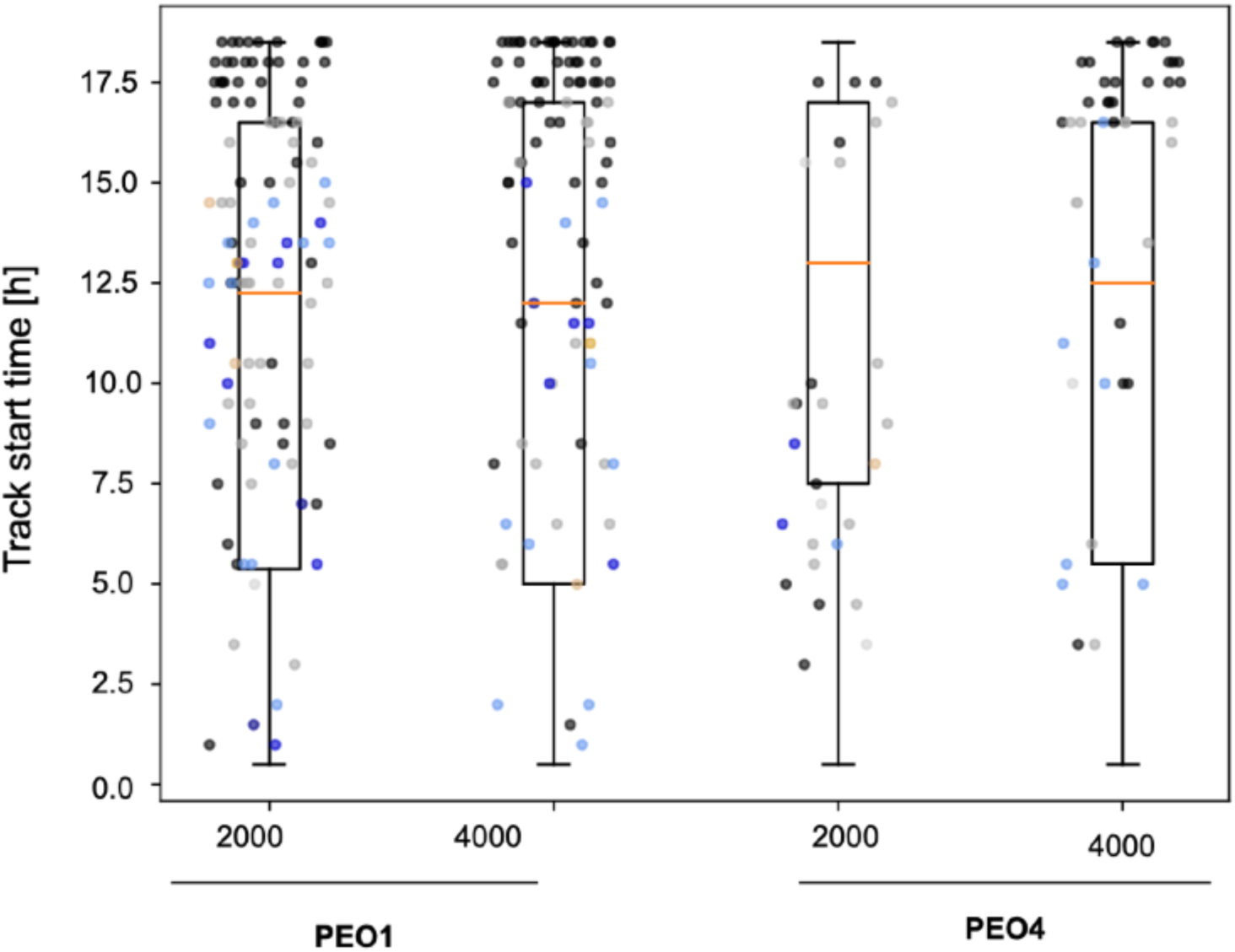
Analysis of the distribution of start times according to cell line and initial cell density. Each dot represents a daughter cell and the colour identify the corresponding track type.

## Discussion

We have here presented an analysis pipeline for the quantification of cell adhesion from microscopy images. This method provides several advantages with respect to the currently available procedures. Indeed, it is characterised by high temporal resolution and by the possibility of distinguishing three different adhesion levels (detached, partially attached and fully attached). The segmentation of individual cells, additionally, enables the study of their individual adhesion dynamic, a unique feature of this approach that yielded interesting insights into the adhesion process and how it changes among cell lines. The pipeline that we have described is also high throughput, largely automated, and its results are operator independent. These features, together with the possibility of changing the instrument used to acquire the images and the segmentation algorithm, suggest that this method could become the new standard for the evaluation of cell adhesion *in vitro*.

Indeed, the population level results that we obtained are largely equivalent to those of more traditional approaches (16) that show PEO1 cells adhering more quickly than PEO4s. While a more extensive analysis of this difference is needed, it might indicate a change in adhesion with OC progression. Earlier stages of the disease, represented by PEO1 cells, might be able to adhere more effectively as this feature would promote cancer dissemination and progression. A more advanced disease (PEO4 cells), on the other hand, might be less dependent on its ability to attach to the healthy tissue and thus divert resources toward development of treatment resistance and improvement of survival. This hypothesis is further supported by the increased variability that we observed for PEO4 cells. Indeed, a more heterogenous population has been associated with increased robustness and improved survival in changing environments (21). Traditional population level measurements, however, are unable to quantify this property as they focus on the difference between populations, rather than among individuals.

Another result presented in this work is the limited effect of changing the initial cell density on the result. This suggests that, in the tested conditions, space availability at the bottom of the plate was not a limiting factor. While unexpected, given the 4-fold change in population size that we considered, this data demonstrates the robustness and accuracy of our approach on a wide range of population sizes, thus further increasing its potential scope of application.

The feature that sets the proposed method apart from the available alternatives, however, is the possibility of analysing adhesion dynamics at the single-cell level. Nine different adhesion dynamics, varying remarkably both in length and complexity, were identified. While the prevalence of each track type was largely conserved across the different tested conditions, full attachment was more common for PEO1 cells, in agreement with the population level results. Track length tended to be proportional to complexity, with cells attaining full adhesion, or following a non-monotonal path generally characterised by longer tracks. A non-negligible variability was however observed. This was particularly obvious for PEO4 cells, that tended to be characterised by longer tracks and a reduced difference between the length of tracks with different dynamics.

Another feature of the tracking algorithm used in this work is its ability to reconstruct the lineage tree for each tracked cell. This enabled us to identify several division events and connect them to specific adhesion dynamics. Indeed, despite having a reported doubling time of 37 and 36-46 h respectively (15), both PEO1 and PEO4 cells underwent division as early as 1 h after the beginning of the experiment (Figures 4 and 9). This proliferation was not associated to a significant growth ratio (Figures 4a and 9a), but the fraction of dividing cells was higher for PEO1 cells, and it seemed to correlate with the initial cell density (Figure 4b). This phenomenon is coherent with a partial preservation of the contact inhibition of proliferation in PEO1 cells. Indeed, while this is a mechanism mostly assumed to be disrupted in malignant cells, recent publications suggest that some cancer cells might still maintain sensitivity to contact inhibition (22). The loss of proliferation inhibition with a larger initial cell population observed for PEO4 cells could be an indication of further disruption of the mechanisms that help maintain a stable cell population as the disease progresses. Coupling this result with the longer track lengths (Figure 3), suggests that PEO4 cells might be more effective in establishing a culture, with cells living longer and thus reducing the need for a high number of cell divisions.

**Figure 9:**
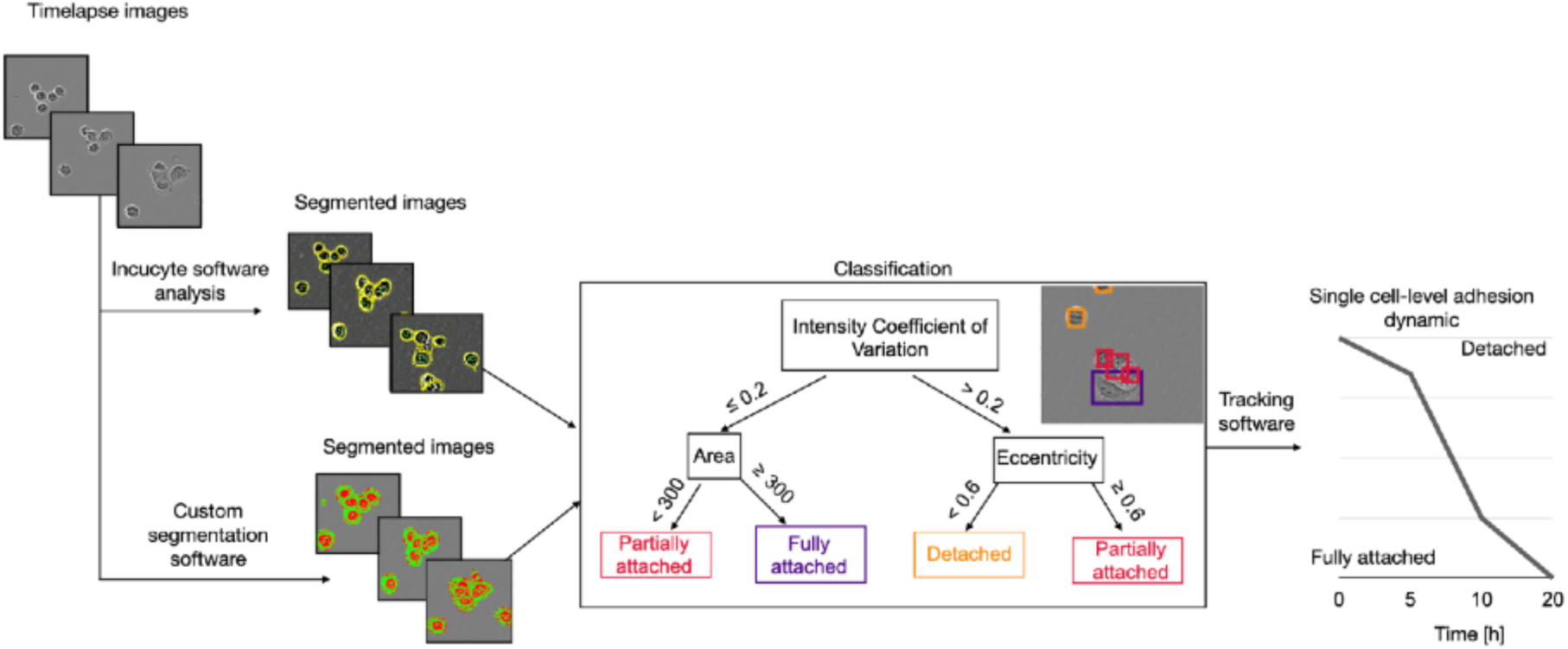
Flowchart providing an overview of the analysis. Timelapse images of PEO1 and PEO4 cells during the adhesion process were segmented using IncuCyte S3 Live Cell Analysis system proprietary software. The segmented cells were then classified according to their adhesion status using morphological and greyscale intensity features. A tracking algorithm was then applied to match individual cells across consecutive images and retrieve their individual adhesion dynamic. The same analysis was then repeated with a custom-made segmentation software (18) to test the dependency of the results on the segmentation software.

An analysis of the adhesion dynamics most likely to be associated with division events (Figures 4d and 7c) highlighted how full attachment is not required for cells to double and that a looser more dynamic attachment might favour early division. Indeed, while disassembly of adhesion complexes and cell rounding has been consistently observed before division to make space for the mitotic spindle and facilitate its alignment (23, 24), full attachment to the substrate has generally been considered as the starting point for this process. The results presented in this work (Figure 4c) paint a more complex picture, with adhesion and proliferation occurring simultaneously and at different rates in different cell sub-populations. While the change in overall cell number was limited (Figures 4a and 7a), no division was expected to occur as the two cell lines used in this work have an average doubling time that is almost twice as long as the experiment length. This raises questions on the accuracy of seeding density as a normaliser between different conditions, especially for cells with short doubling time and encourages the development and use of experimental techniques able to capture the complexity and variability of biological processes.

### Conclusions

In this work, we have presented a novel approach for the quantification of cell adhesion *in vitro*. This method addresses the major limitations of the standard approach for the evaluation of adhesion, high processivity and lack of standardisation, while yielding more complete information. Indeed, it enables a notable increase in the temporal resolution of the analysis, while providing a breakdown of three different adhesion stages and enabling the study of adhesion at the single cell level.

The independence of the results from the segmentation algorithm, demonstrated by repeating the analysis using two different methods, greatly expands the potential and scope of the proposed approach, making it independent of the live cell monitoring instrument available and potentially applicable to standard microscopy setups.

Finally, the results obtained with this method shed new light on the complexity and variability of cell adhesion and could be the base for further investigations into the role of doubling time and other cell- and environment-specific factors in this process.

## Methods

### Overview of the analysis pipeline

Figure 9 provides an overview of the proposed method. Images of live OC cells adhering to the bottom of the plate were segmented to retrieve the outline of each cell. Two different algorithms were used at this stage to test the dependency of the results on the proprietary software of the IncuCyte S3 Live Cell Analysis system.

Specific morphological and intensity features were then used to classify each cell according to its adhesion status (detached, partially attached or fully attached). This enabled the study of the prevalence of each adhesion class and how it changed over time. This analysis was repeated changing the initial cell density, to determine if space availability at the bottom of the plate would influence cell adhesion.

The btrack tracking software (17) was then used to match individual cells across consecutive images and retrieve their specific adhesion dynamic. This increase in resolution is one of the main features of this analysis pipeline and provided novel and interesting insights into cell adhesion.

### Cell culturing and adhesion experiments

OC cell lines PEO1 and PEO4 were kindly donated by Prof. Deborah Marsh (University of Technology Sydney, NSW). They were maintained in RPMI medium (Thermo Fisher, USA) supplemented with 10% FBS (Sigma-Aldrich, USA), 1% Pen-Strep (Sigma-Aldrich, USA) and 1% GlutaMAX (Thermo Fisher, USA) in standard cell culture conditions (37°C and 5% CO_2_). For the adhesion experiment, three different densities (1000, 2000 or 4000 cells/well) were considered for each cell line. Cells were seeded in a 96-well plate (i.e., Nunclon Delta-treated, Flat-Bottom, 6 wells/condition, 100 µl/well) and immediately transferred to the IncuCyte S3 Live Cell Analysis System (Sartorius, Germany). Three images for each well were acquired with a magnification of 10x every 30 minutes for a total of 20 hours.

### Incucyte software segmentation and adhesion status determination

The IncuCyte proprietary analysis software was used to analyse the images and retrieve the cells’ outlines. All the segmentation parameters were left at their default value. Images were then exported in tif format. The segmentation was completed by filling holes with area < 2000 px. Objects comprising 50 px or less were considered to be debris and disregarded. A custom-made Python (v 3.9) script was used to determine the adhesion status of each segmented cell (Figure 9). It relies on three parameters: eccentricity (E), area (A) and coefficient of variation of the intensity (CV). Detached cells are round and have high contrast, as such cells with an E < 0.6 and CV > 0.2 were classified as detached. Conversely, fully attached cells were identified by their larger area (A ≥ 300 px) and lower contrast (CV ≤ 0.2). All the other cells were assigned to the intermediate partially attached group (Figure 4).

### Custom-made segmentation algorithm

The custom-made segmentation algorithm used in this work was initially described in (18). It was programmed with MATLAB 2021b Image Processing Toolbox^TM^. Cells on phase contrast image stacks were divided into sections based on relative pixel intensity compared to image background. Segmented dark areas (green) or light areas (red) were correlated to different parts of the cell that were, respectively, adherent and non-adherent to the substrate. The total number of pixels in the dark areas per field of view was used to calculate the estimated cell attachment area.

For the analysis here presented only the dark areas (green channel) were used and, as with the previous algorithm, objects with area <50 px were disregarded. No modification to the adhesion classification routine or the tracking software was implemented.

The similarity of the two segmentations was evaluated through the Jaccard index (Eq. 1). This metric was computed independently for each image and then grouped according to the experimental condition (cell line and initial cell density).

### Global adhesion analysis

Population-level adhesion was quantified by computing, for each time-point and experimental condition, average and standard deviation of the prevalence of each adhesion status. The number of cells in the first image of each series was used as normaliser.

The Bland Altman plot was used to compare the results of the global analyses obtained with the two segmentations. This graph plots the difference between the results of the two methods as a function of their average. Horizontal lines identify the average and 95% confidence interval of the difference.

### Tracking software

The Bayesian tracking software btrack (17) was used to match individual cells across images and retrieve their adhesion dynamic. It relies on a Bayesian belief matrix and a cell motion model construct tracklets, that describe the position of each cell up until cell division. Multiple hypothesis testing is then performed to assemble the tracklets into tracks and retrieve each cell’s lineage tree (17). For some of the image stacks (Table 3) the optimisation failed. Changing the optimisation parameters might have impacted the comparison between different conditions, as they directly relate to the model of cell movement. As such, we decided to exclude these images from this analysis. As no data were available for the condition PEO1 cells, 1000 cells/well using the custom segmentation method, we decided to exclude this cell density from the analysis relying on this approach.

**Table 3:**
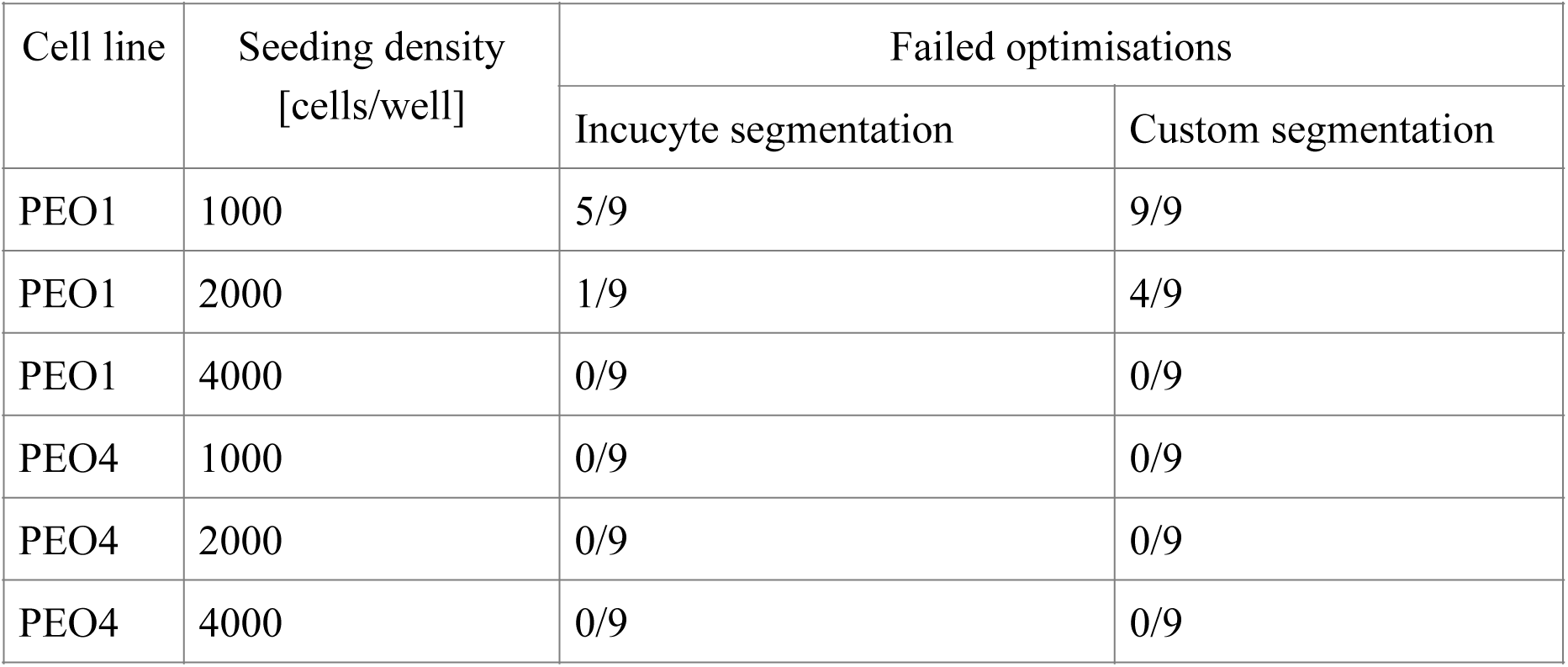
Summary of the failed btrack optimisations by experimental condition and segmentation method.

### Single-cell level adhesion analysis

The tracks were analysed to determine how each cell’s adhesion status would change throughout the experiment. The same classification procedure described in the previous section (Figure 9b) was applied, and the resulting adhesion tracks were smoothed applying a moving average with window size equal to 3. This procedure was used to avoid rapid short switches in the adhesion status, which would be likely caused by the cell’s properties being close to the classification thresholds. This step was not applied to tracks shorter than 4 h as per the definition of moving average.

This analysis identified nine different adhesion dynamics (Table 1). Individual tracks were sorted automatically according to the following procedure. Short tracks were identified by their length (< 2 h). The range of the variation then was calculated. If it was less than 1/6 of the available dynamic range the track was classified as flat. The distinction between flat detached and partially attached was made according to the corresponding adhesion status. The remaining tracks were then divided between monotone and non-monotone. For monotone tracks the last recorded adhesion status was compared to the first one. This enabled the distinction between detaching and attaching cells. The last adhesion status was also used to distinguish between monotonal partially attached and monotonal fully attached. A similar approach was used to identify non-monotonal partially attached tracks. All the remaining dynamics were non-monotonal with full attachment. The number of adhesion events was evaluated as the number of continuous stretches of the track classified as being fully attached (2 in Figure 2d).

The prevalence of each adhesion dynamic was determined as the average fraction of tracks in each class following the exclusion of the short tracks. This was necessary due to the high prevalence of short tracks with late start time (Figure 5) which would have overshadowed the dynamics observed at earlier timepoints.

### Analysis of the doubling events

The optimisation step in btrack enables the reconstruction of the lineage trees and thus the identification of division events and of the cells involved in this process. In our analysis we focussed both on the dividing cells, and on the cells that were generated during the experiment. For the former we determined their prevalence within the population (number of cells determined to have undergone division divided by the total number of tracked cells), the adhesion status immediately prior to cell doubling and adhesion dynamic. The adhesion dynamic was evaluated also for the newly born cells, together with the distribution of the start times for their track.

For the subset of cells attaining full adhesion the cumulative distribution of the time interval between full attachment and division was also calculated.

The absolute value of the difference between the fraction of dividing cells obtained with the either segmentation method was also computed to provide an indication of the disparity of their results.

### Statistical Analysis

Statistical analysis was conducted using the Kruskal-Wallis test, a non-parametric test which compares the median of two populations to determine whether or not they are equal. A p value of 0.05 was considered as threshold for significance.

## Supplementary Figures

**Supplementary Figure 1:**
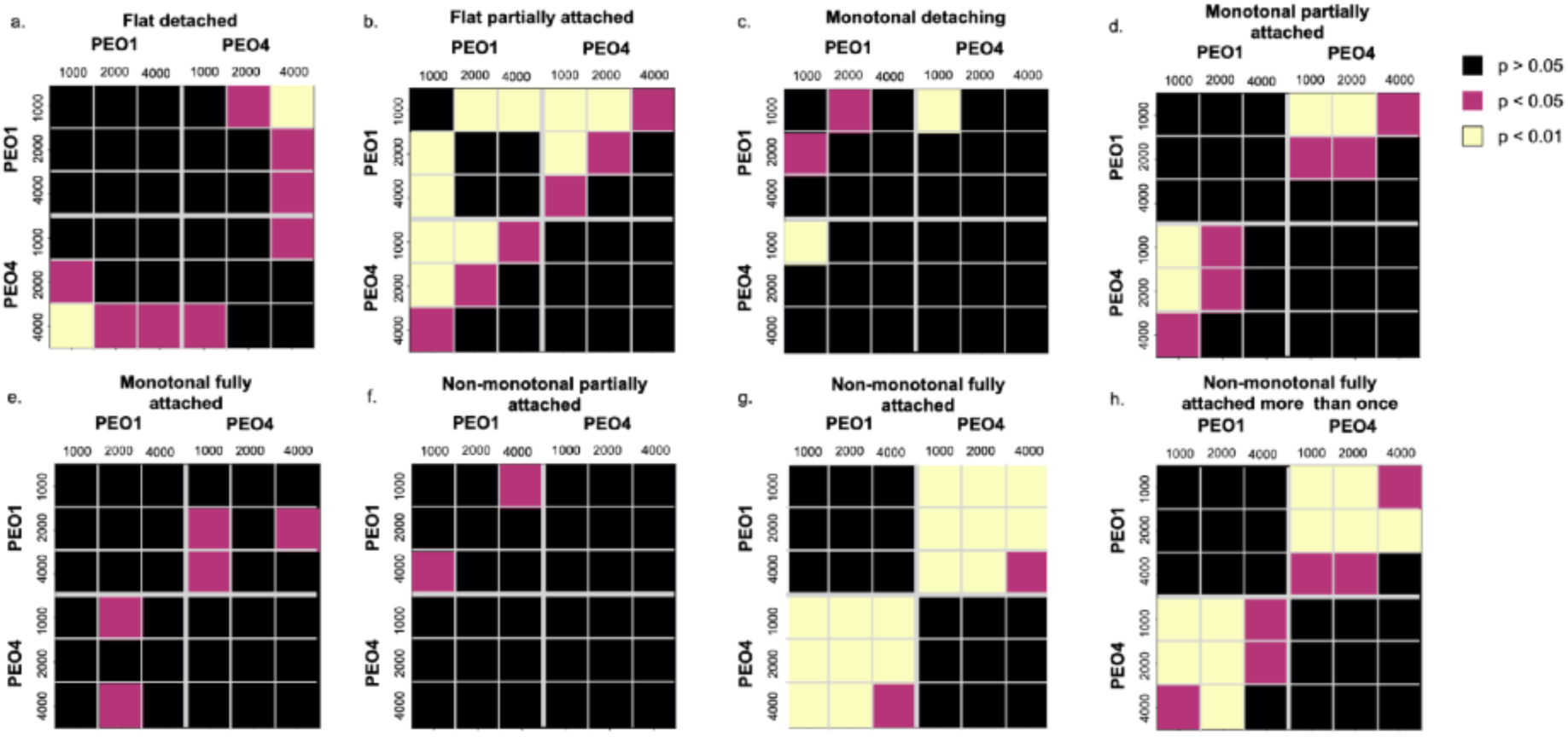
Statistical analysis of the prevalence of each adhesion dynamic. The Kruskal-Wallis test was used to compare each condition and a p value of 0.05 was chosen as threshold for significance.

**Supplementary Figure 2:**
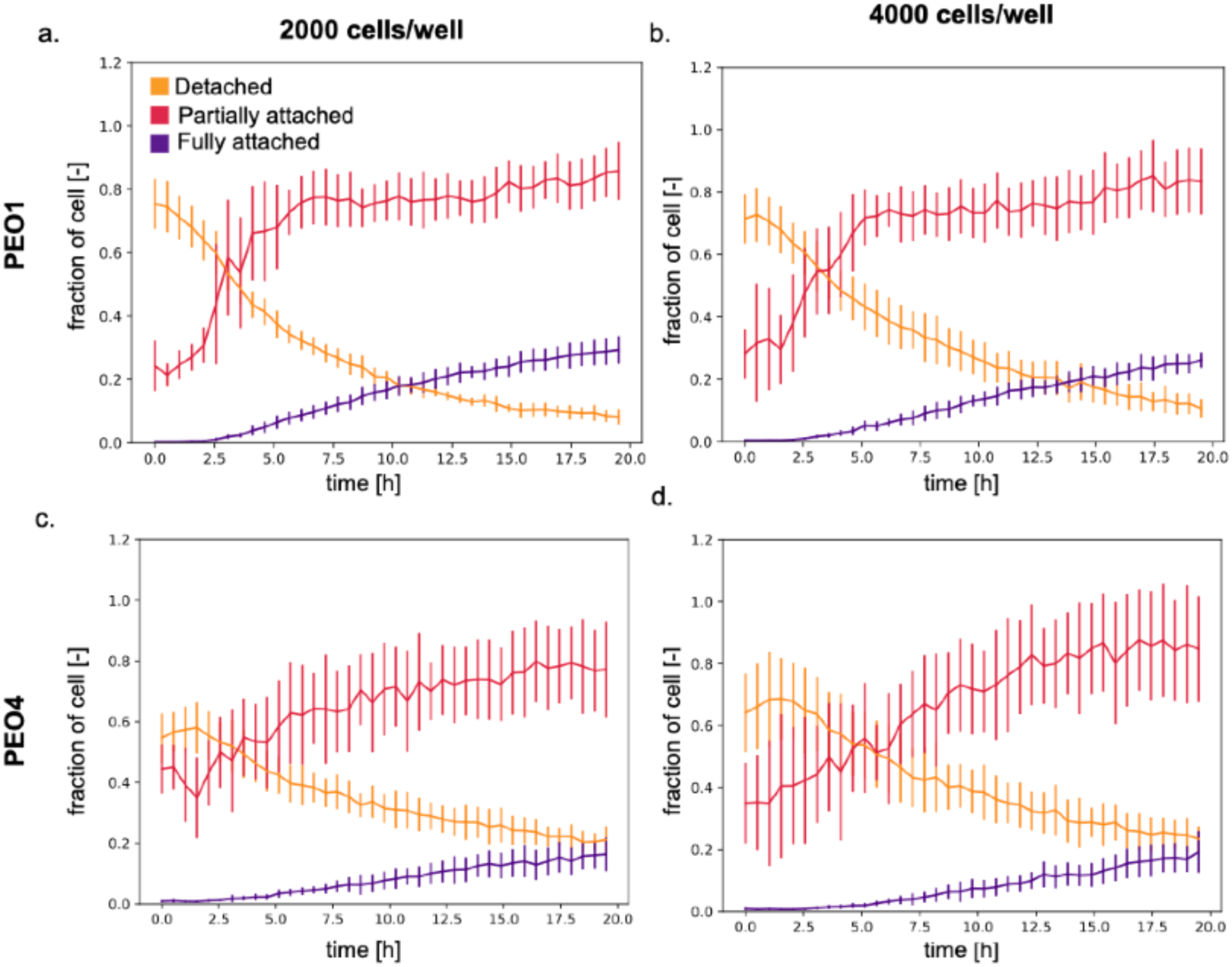
Global adhesion analysis for the custom segmentation algorithm. a. Average and standard deviation of the fraction of cells in each adhesion status measured for PEO1 cells with an initial cell density of 2000 cells/well. b. is the same as in a. but with starting populations of 4000 cells/well. C. and d.show the same analysis but for PEO4 cells. In all cases 3 wells/condition and 3 images/well were considered.

**Supplementary Figure 3:**
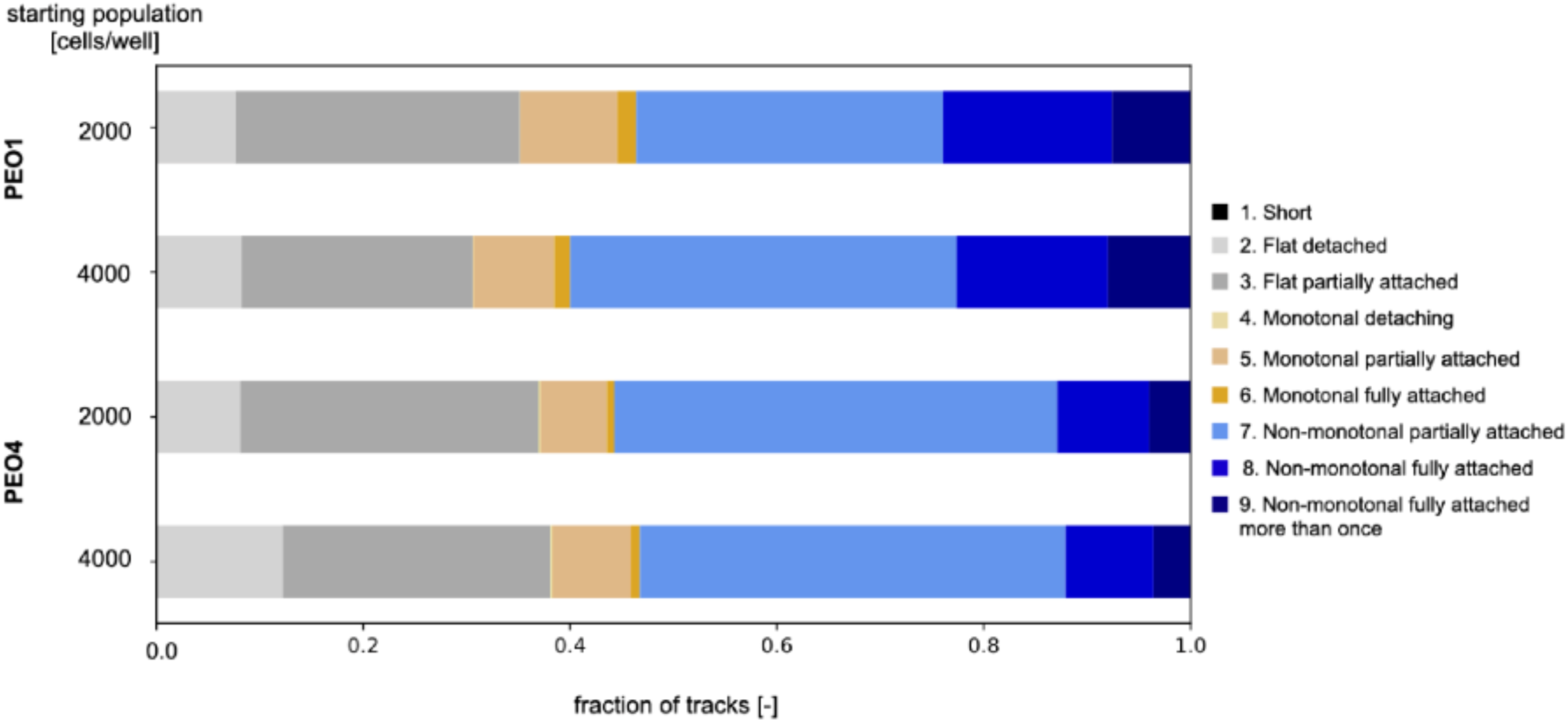
Prevalence of each adhesion type retrieved with the custom segmentation algorithm. As in Figure 3e short tracks were excluded.

**Supplementary Figure 4:**
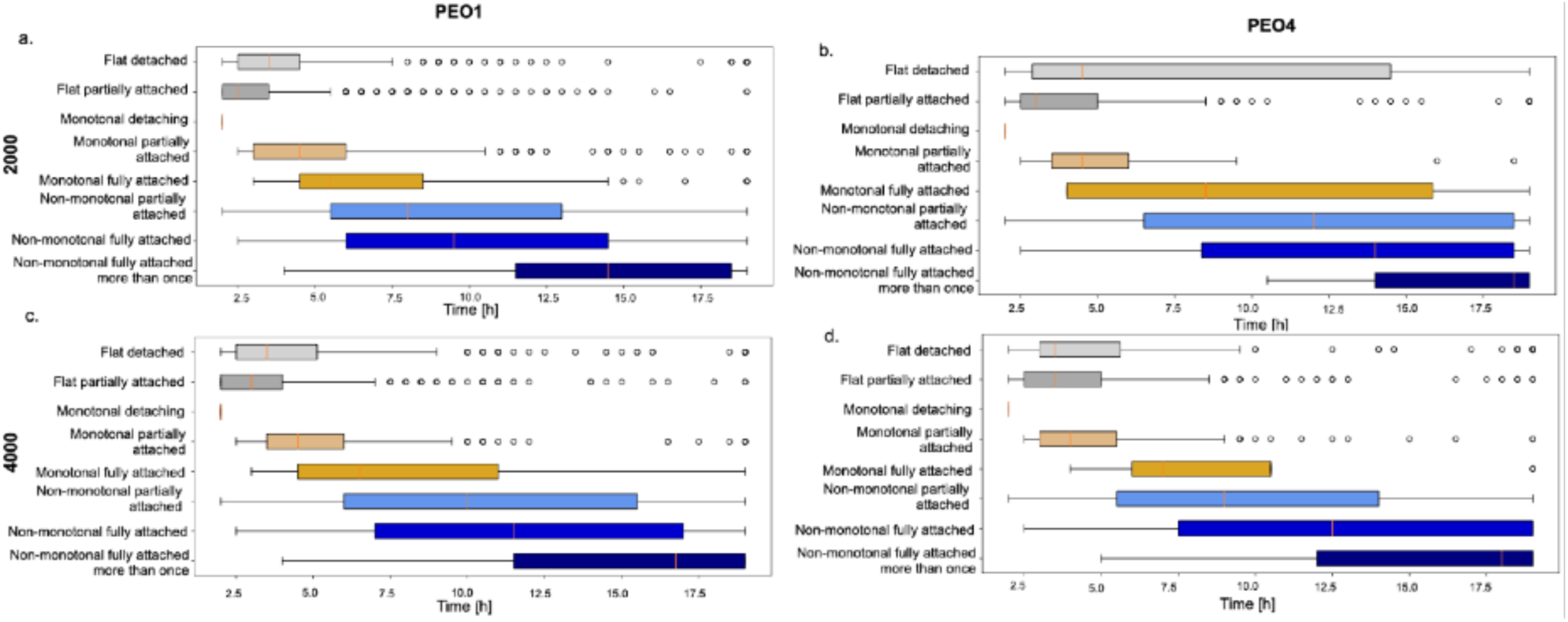
Study of the distribution of track length as a function of cell line, initial cell density and adhesion dynamic. Segmentations obtained with the custom algorithm.

## Notes

### Competing Interest Statement

The authors have declared no competing interest.

