## Supplementary figures and images for "A novel approach for the quantification of single-cell adhesion dynamics from microscopy images"

### Supplementary Figure 1

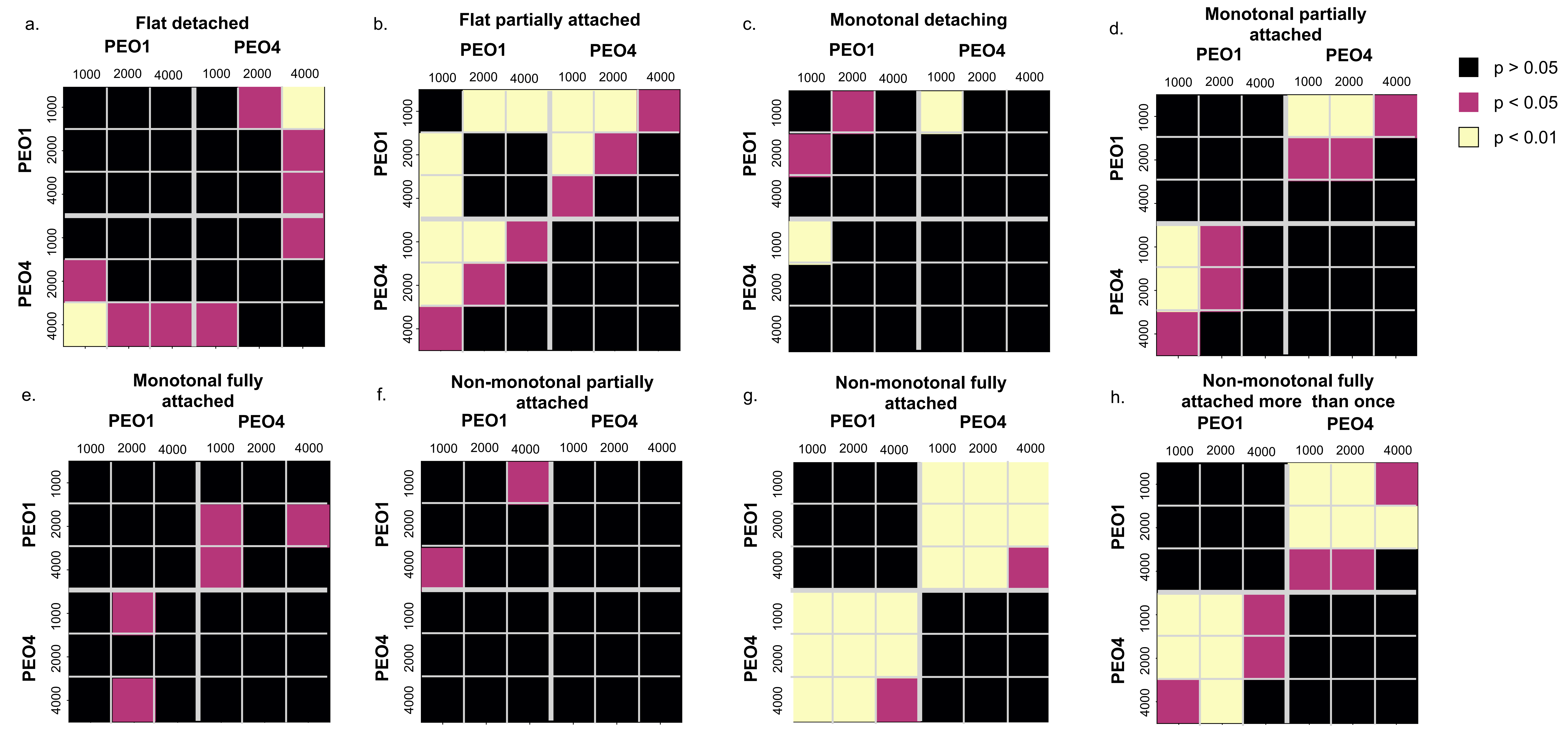

### Supplementary Figure 2

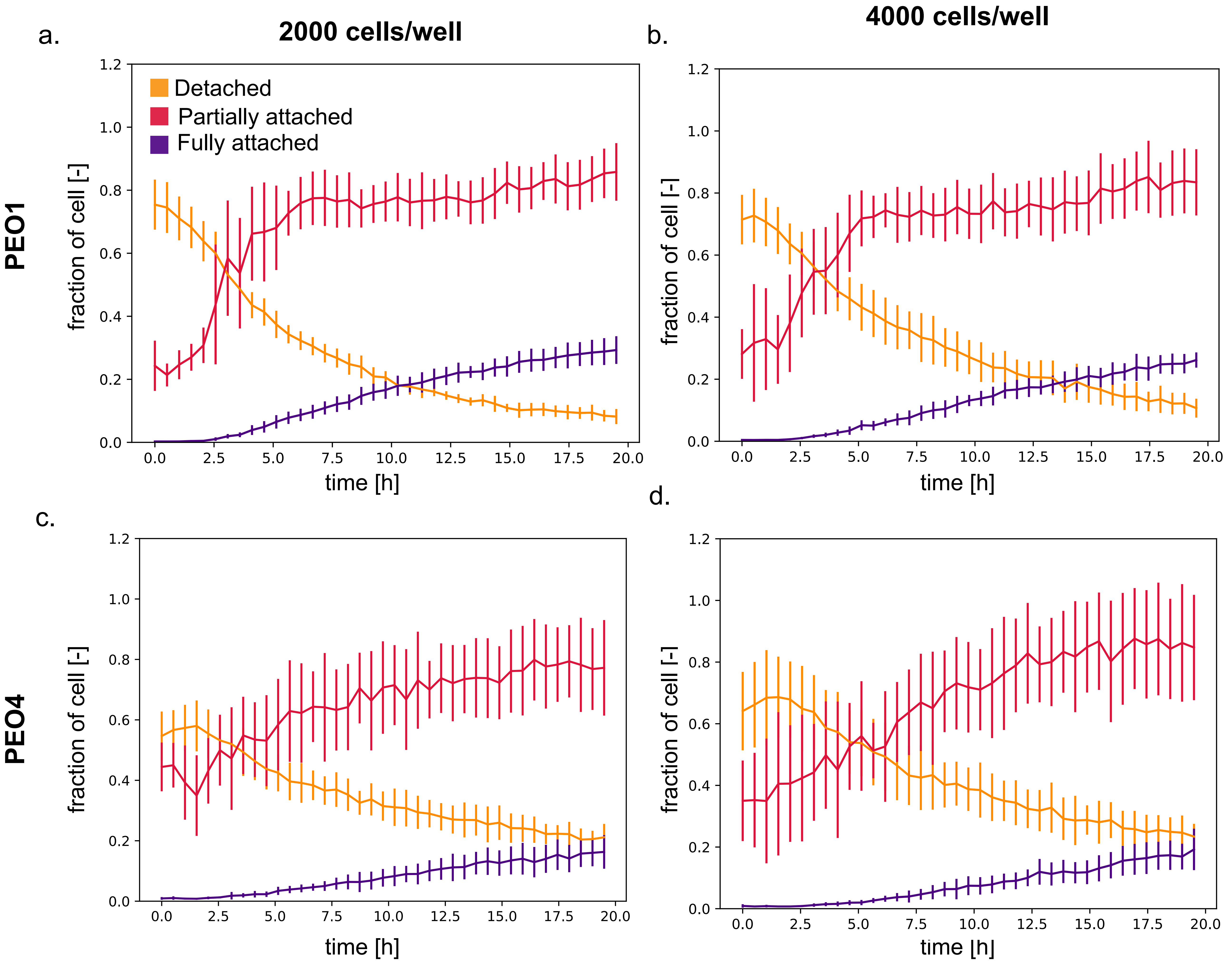

### Supplementary Figure 3

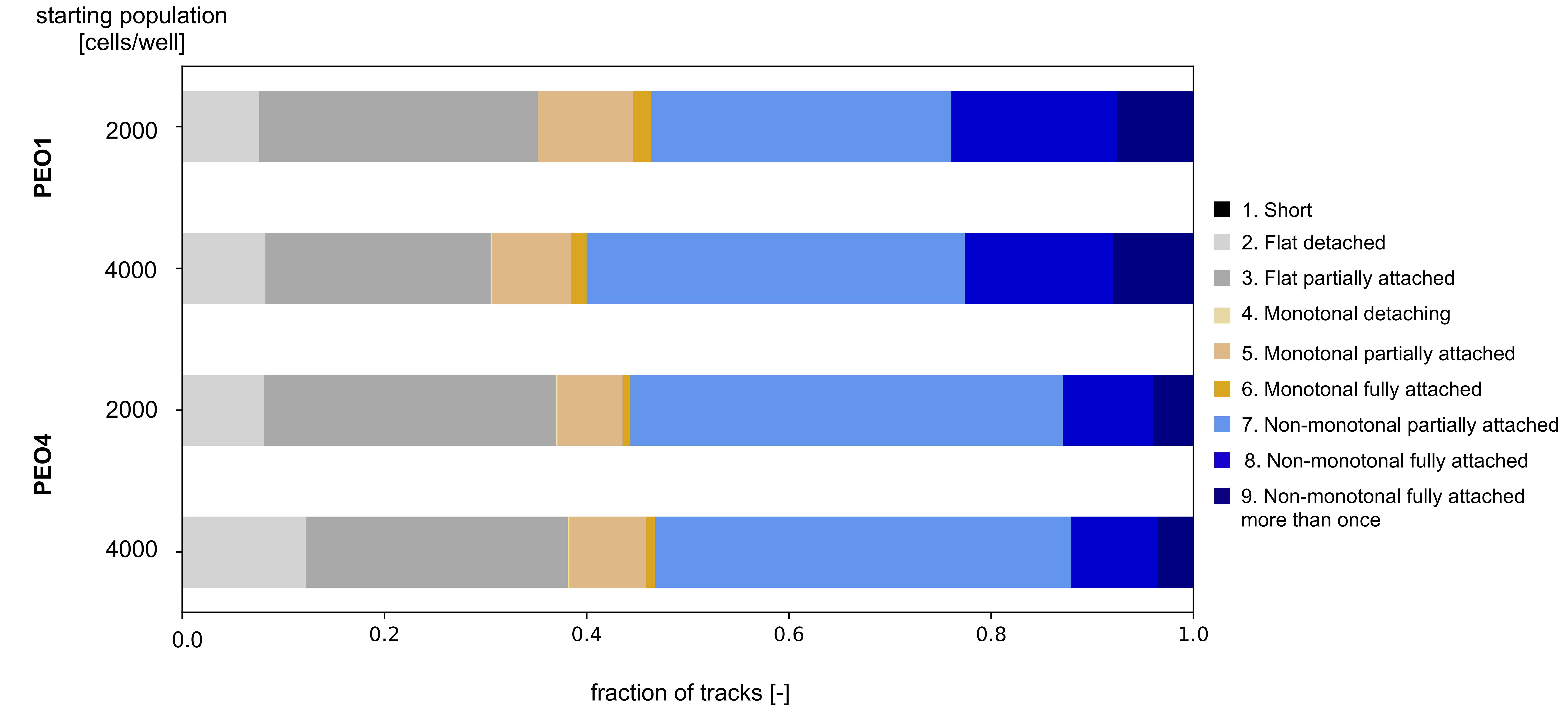

### Supplementary Figure 4

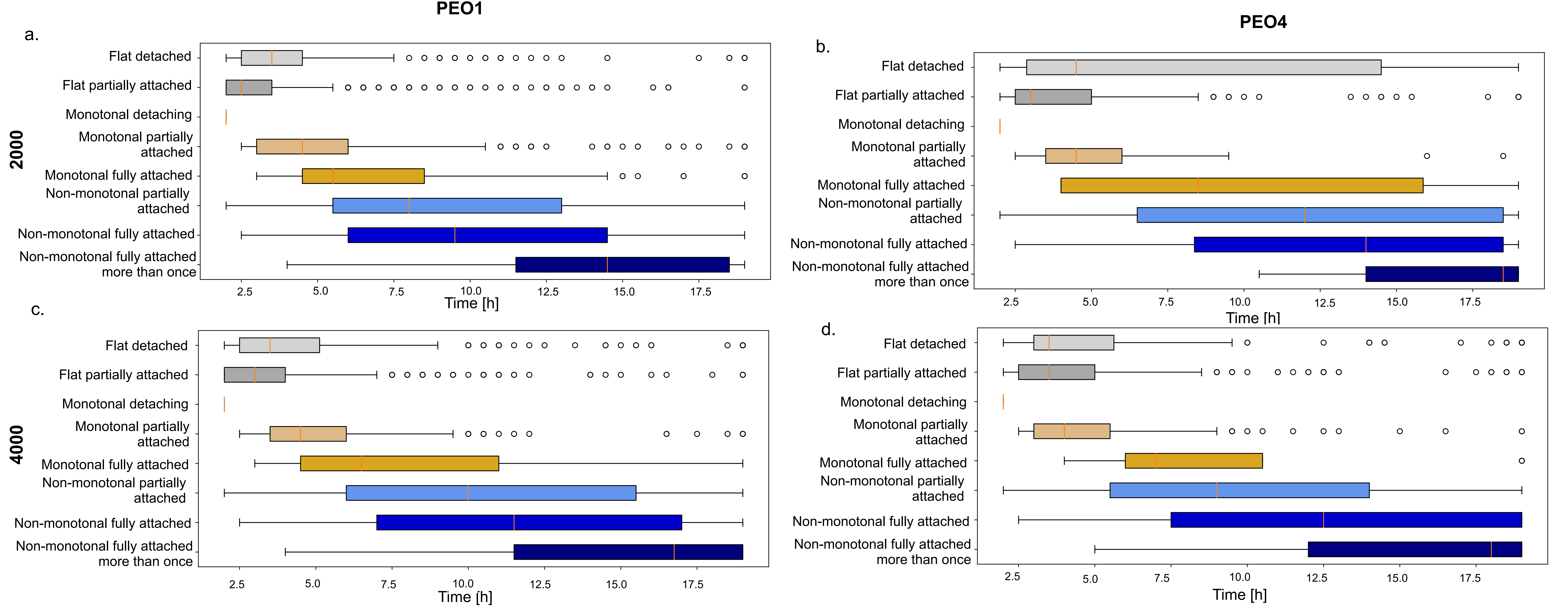
